# Effects of BCG vaccination on donor unrestricted T cells in humans

**DOI:** 10.1101/2021.04.29.441927

**Authors:** Anele Gela, Melissa Murphy, Kate Hadley, Willem A. Hanekom, W. Henry Boom, John L. Johnson, Daniel F. Hoft, Simone A. Joosten, Tom H.M. Ottenhoff, Sara Suliman, D. Branch Moody, David M. Lewinsohn, Mark Hatherill, Chetan Seshadri, Elisa Nemes, Thomas J. Scriba, The Delayed BCG Study Team

## Abstract

Antigen classes other than proteins can be presented to T cells by near-monomorphic antigen-presenting molecules such as CD1, MR1, and butyrophilin 3A1. We sought to define the roles of donor unrestricted T (DURT) cells, including MR1-reactive MAIT cells, CD1b-reactive glucose monomycolate (GMM)-specific T cells, CD1d-reactive NKT cells, and γδ T cells, in vaccination against *Mycobacterium tuberculosis*. We characterized DURT cells following primary bacille Calmette-Guerin (BCG) vaccination in infants or BCG-revaccination in adults. BCG (re)vaccination did not modulate peripheral blood frequencies, T cell activation or memory profiles of MAIT cells, CD1b-restricted GMM-specific and germline-encoded mycolyl-reactive (GEM) cells or CD1d- restricted NKT cells. By contrast, BCG vaccination was associated with increased frequencies of γδ T cells as well as a novel subset of IFN-γ-expressing CD4^+^ T cells with a CD26^+^CD161^+^TRAV1-2^−^ phenotype in infants. More studies are required to understand the full potential of DURT cells in new TB vaccine strategies.

## Introduction

Current approaches to develop vaccines against tuberculosis (TB), one of the top 10 causes of death globally, are based primarily on the induction of conventional, major-histocompatibility complex (MHC)-restricted, Th1 cytokine-producing T cells to protein antigens. While proteins constitute the majority of known antigens recognized by αβ T cell receptor (TCR)-bearing cells, other chemical classes including lipids, small-molecule metabolites and specially modified peptides are also antigenic and can be presented to T cells by the non-polymorphic molecules, CD1, MR1, and butyrophilin 3A1, respectively (1–3). Unlike MHC-restricted T cells, these T cell subsets generally bear T cell receptors that show limited diversity, and are therefore referred to as donor-unrestricted T (DURT) cells (3). DURT cells are activated through their TCR in a manner that is unrestricted by donor origin: response is mediated by antigen presentation molecules of limited polymorphism, such that DURT responses can be shared among genetically diverse individuals (1). This nature of antigen recognition by DURT cells is of particular interest for vaccine development, since immunogens could be designed to elicit universal, population-wide T cell responses, irrespective of host MHC-encoded genetic factors.

Studies in mouse and non-human primate models have demonstrated the importance of T cells in conferring protection against challenge with *Mycobacterium tuberculosis* (M.tb) (4–6). However, it remains unclear which and how many mycobacteria-derived antigens should be targeted by protective T cell responses, and which T cell functional, homing and memory attributes are required for protection (7). Given the complexity of host-pathogen interactions, TB vaccination strategies that exploit immunological diversity and target additional arms of the immune system, including DURT cells, such as the CD1d-restricted NKT cells, CD1b-restricted glucose monomycolate (GMM)-specific T cells and germline-encoded mycolyl-reactive (GEM) cells, γδ T cells, and MR1-restricted MAIT cells, should be explored (1).

Current studies provide extensive evidence that DURT subtypes are activated during natural M.tb infection in humans and non-human primates. Analysis of the T cell response during human M.tb infection indicates that infected individuals have increased CD1-restricted T cell responses to lipid antigens compared to M.tb naïve controls (8–10). A recent study in non-human primates also reported that glucose monomycolate (GMM)-specific CD1c-restricted T cells expanded after intravenous bacille Calmette-Guerin (BCG) administration (11). Vaccination of guinea pigs with mycobacterial lipids formulated in liposomes was also associated with a reduction in the number of lesions, severity of pathology, and reduction of bacterial load upon challenge with M.tb (12).

Activation of MR1-restricted MAIT cells, which recognize vitamin B metabolites presented by the monomorphic MHC class 1-related molecule (MR1), has been shown in response to a variety of bacteria including BCG, *Francisella tularensis*, *Klebsiella pneumonia*, and M.tb (13). Despite previous evidence that MAIT cells can respond to M.tb infection, there is conflicting data on the importance of MAIT cells in controlling infection. In the murine model, endogenous MAIT cell responses have been implicated in early innate responses to M.tb infection (14), but more recently additional studies suggest they are less important for adaptive host defenses against M.tb infection (15–17). Moreover, early MAIT cell responses impeded the priming and induction of conventional peptide-specific CD4 T cell responses (15).

Butyrophilin 3A1 facilitates recognition of phosphoantigens, small molecules like isopentyl pyrophosphate, by γδ T cells, which rapidly respond to pathogen infection in mucosal tissues, and are induced by both M.tb infection and BCG vaccination in humans and non-human primates (18–22). Importantly, induction of Vγ2Vδ2 T cells specific for the phosphoantigen, (E)-4-hydroxy-3-methyl-but-2-enyl pyrophosphate (HMBPP), in non-human primates was associated with increased Th1-like Vγ2Vδ2 T cells in the airways, facilitated earlier recruitment of conventional Th1 cytokine-expressing CD4 and CD8 T cell to the lungs, and was associated with containment of M.tb after pulmonary challenge (20).

Another feature of DURT cells that makes them attractive as vaccine targets is their inherent immediate effector function, such as secretion of inflammatory cytokines and cytotoxic molecules upon recognition of microbial antigens (23). Immediate cytokine secretion allows DURT cells to modulate the antimicrobial function of other cells before recruitment or induction of conventional MHC-restricted T cells has occurred. For example, MAIT cell secretion of CCL4 mediates recruitment of NK cells, monocytes, and other inflammatory cells to infected tissues (24). Similarly, immediate or early secretion of pro-inflammatory cytokines that can facilitate the development of adaptive immunity has been described for NK cells (25), γδ T cells (26), and NKT cells (27). The relatively high baseline precursor frequency in blood and tissues of DURT cells allows them to recognize antigens and respond in large numbers without requiring extensive clonal expansion, implying that they can act simultaneously with innate cells as sensors of infection or immune dysregulation. Despite these attributes, it remains unknown whether DURT cells possess immunological memory such that they can be selectively expanded by vaccination to provide long-term, antigen-specific protective immunity.

In this study, we sought to determine if a live attenuated vaccine, BCG, modulates frequencies, phenotypes or functions of DURT cells in humans.

## Results

### Participant enrollment and demographic data

We enrolled two cohorts from communities residing in a TB endemic setting, in South Africa. The infant cohort comprised two groups of healthy 9-week-old infants, born to HIV negative mothers, who either received routine BCG at birth or in whom BCG vaccination was delayed until after blood collection at 9 weeks. There were no significant differences in the gender and ethinicty distribution between the vaccinated and the delayed BCG arm. The adult cohort included tuberculin skin test (TST)-positive, HIV-negative adults who received BCG revaccination after pre-treatment with isoniazid as part of the TBRU adult BCG revaccination trial (28, 29). Peripheral blood samples were analyzed before revaccination and on days 21, 35, and 365 post-vaccination. The demographics of the study populations are summarized in Table 1.

**Table 1:** Demographic characteristics of participants.

| Variable | Infants | Adults |
| --- | --- | --- |

|  | No BCG (N = 25) | BCG (N = 50) | (N = 25) |
| --- | --- | --- | --- |
| BCG (re)vaccination | No | Yes | Yes |
| Median age | 9 weeks | 9 weeks | 24 years |
| Range | (8-11) | (8-11) | (19-39) |
| Sex, n (%) |  |  |  |
| Male | 13 (52) | 25 (50) | 7 (28) |
| Female | 12 (48) | 25 (50) | 18 (72) |
| Ethnicity, n (%) |  |  |  |
| African | 4 (16) | 13 (26) | 5 (20) |
| Caucasian | 0 (0) | 0 (0) | 1 (4) |
| Mixed ancestry | 21 (84) | 37 (74) | 19 (76) |

### DURT cell subset characterization by flow cytometry

We quantified CD1b-, CD1d-, MR1-restricted, and γδ T cells in peripheral blood using a combination of tetramers and monoclonal antibody staining against phenotypic markers. DURT cell subsets were defined by flow cytometry as MR1-5-OP-RU^+^TRAV1-2^+^ (or CD26^+^CD161^+^) MAIT cells, CD1d-PBS57^+^ NKT cells, γδ-T cells, and CD1b-GMM^+^CD4^+^TRAV1-2^+^ GEM cells (Fig. 1).

**Figure 1.**
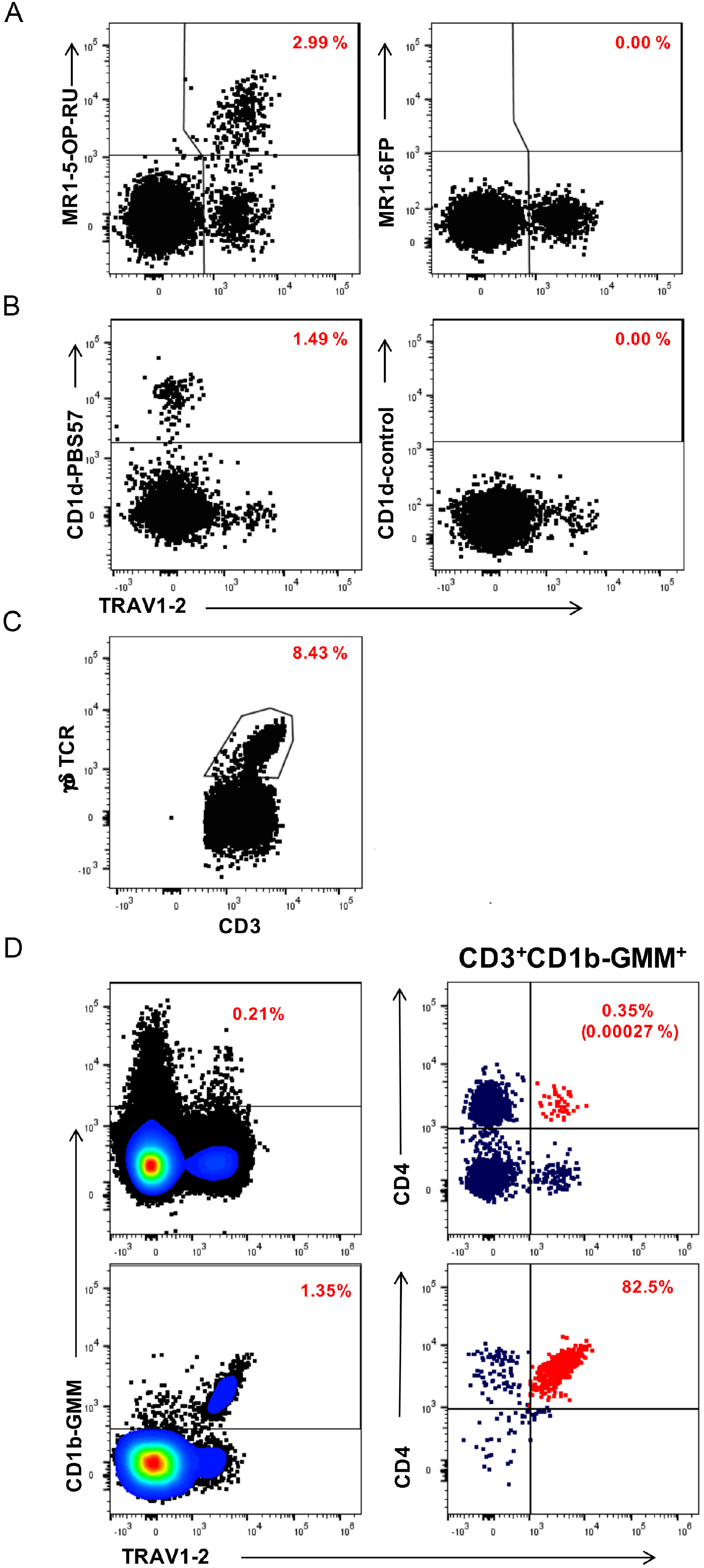
Representative flow cytometry plots demonstrating detection of DURT cell populations in a typical infant. Plots represent live, CD3+ T cells co-stained with the indicated antibody-conjugate on the X-axis and the indicated tetramer reagent or antibody-conjugate on the Y-axis. (A) MR1-restricted MAIT cells (MR1-5-OP-RU tetramer on the left and MR1-6FP tetramer as negative control on the right). (B) CD1d-restricted NKT cells (CD1d-PBS57 tetramer on the left and unloaded [empty] CD1d tetramer as negative control on the right). (C) γδ T cells, defined as CD3 T cells that stained positive with anti-pan γδ TCR antibody. (D) Definition of CD1b-restricted glucose monomycolate (GMM)-specific T cells and germline-encoded mycolyl-reactive (GEM) cells in an infant sample (top plots) or a PBMC sample that was spiked with GEM T cell clone cells (bottom plots). GMM-specific T cells (left plots) were defined as CD1b-GMM tetramer^+^ T cells and GEM cells were defined as CD1b-GMM tetramer^+^ T cells that co-expressed CD4 and the TCR variable chain TRAV1-2 (red dots in right plots). Red numbers in all plots denote the percentages of T cells in that plot that stain positive for each population-defining marker. The red numbers in parentheses in the top, right plot in panel D denote the percentage of GEM cells out of all CD3+ T cells.

### Frequencies of γδ T cell, but not other DURT subsets, increase after BCG vaccination in infants

To determine if BCG vaccination alters DURT cell abundance in peripheral blood, we measured frequencies of DURT cell subsets in infants and adults at 35 days post revaccination. Frequencies of the different DURT subsets were generally lower in infants compared to adults, but there was high heterogeneity among the adults, especially for frequencies of MAIT and γδ T cell populations (Fig. 2A-E). Frequencies of MAIT cells, CD1d- restricted NKT cells, CD1b-restricted GMM-specific and germline-encoded mycolyl-reactive (GEM) cells were not significantly modulated by primary BCG vaccination, nor by BCG re-vaccination at all time-points (Fig 2A-D, data not shown for day 21 and 365). However, frequencies of γδ T cells were significantly higher in BCG-vaccinated infants compared to their unvaccinated counterparts (Fig 2E). No difference in γδ T cell frequencies was observed before and after BCG revaccination in adults at any of the post-vaccination time points (Fig 2E, data not shown for day 21 and 365). Infants also had higher frequencies of bulk CD4 T cells than adults, but these were not modulated by BCG in either age group (Fig. 2F). In contrast to these DURT cell subsets, antigen-specific CD4 T cells were significantly modulated by BCG vaccination, as expected for peptide-responsive, MHC-restricted T cells; primary BCG vaccination induced significant increases in BCG-reactive IFNγ+ CD4 T cells in infants, while BCG re-vaccination boosted pre-existing BCG-reactive CD4 T cells in adults (Fig. 2G). Taken together, these results show that with the exception of a significant increase in total γδ T cells in BCG-vaccinated infants, DURT cell frequencies in peripheral blood were not modulated by BCG vaccination.

**Figure 2.**
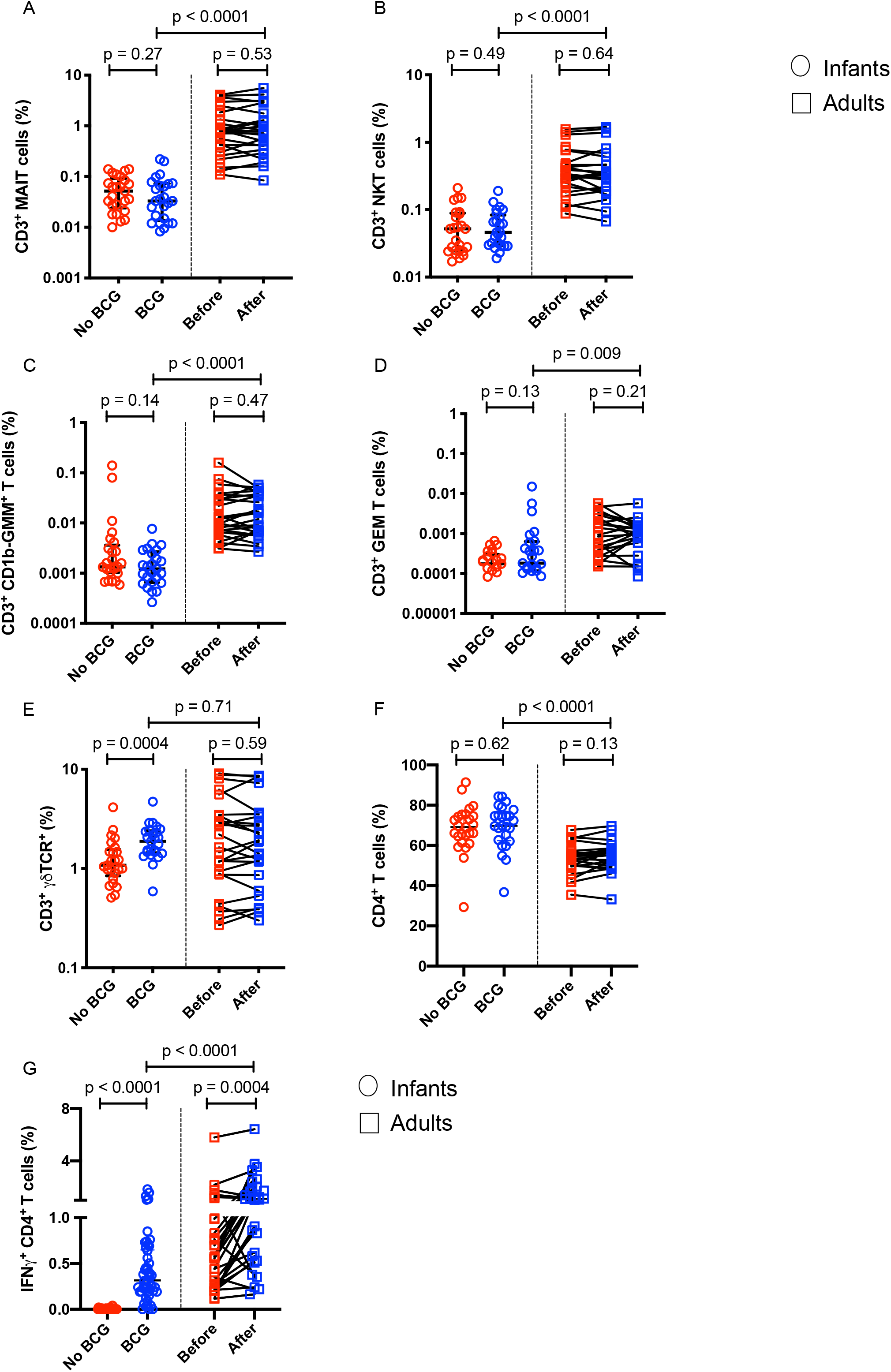
T cell subset frequencies measured by flow cytometry in BCG-vaccinated or unvaccinated infants or before and after BCG revaccination in adults. Peripheral blood frequencies of (A) MR1-5-OP-RU tetramer^+^ MAIT cells, (B) CD1d-PBS57 tetramer^+^ NKT cells, (C) CD1b-GMM tetramer^+^ T cells, (D) CD1b-GMM tetramer^+^ CD4^+^TRAV1-2^+^ GEM cells, (E) γδ T cells and (F) “conventional” CD4^+^ T cells in individual infants (circles) or adults at 35 days post re-vaccination (squares). (G) Peripheral blood frequencies of BCG-reactive “conventional” CD4 T cells expressing IFN-γ in individual infants (circles) or adults (squares). *P* values were computed by Mann-Whitney U test (comparisons between infant groups or between infants and adults) or Wilcoxon signed-rank test (pre- and post-BCG comparisons in adults). p < 0.05 was considered statistically significant.

### BCG vaccination does not modulate the activation status of DURT cells in peripheral blood

We next sought to determine if BCG vaccination results in activation of DURT subsets in peripheral blood, since evidence of T cell activation may indicate that DURT cells sense and respond to the vaccine, or its downstream effects. As expected, BCG-specific IFNγ-expressing CD4 T cells in the BCG vaccinated infants expressed significantly higher levels of the *in-vivo* activation marker, HLA-DR, than bulk CD4 T cells (Fig. 3A). BCG-specific CD4 T cell responses were not detected in unvaccinated infants, precluding analysis of activation status of these CD4 T cells in this group (Fig. 3A). By contrast, HLA-DR expression levels on DURT cell subsets were not different between BCG-vaccinated and unvaccinated infants (Fig. 3B-E). Similarly, levels of HLA-DR expression by DURT cell subsets or bulk CD4 T cells were not different between pre- and post- BCG-revaccination time points in adults (Fig. 4A-E). Taken together, these data suggest that DURT cells in the peripheral blood were not activated by BCG vaccination at the time points that we assessed.

**Figure 3.**
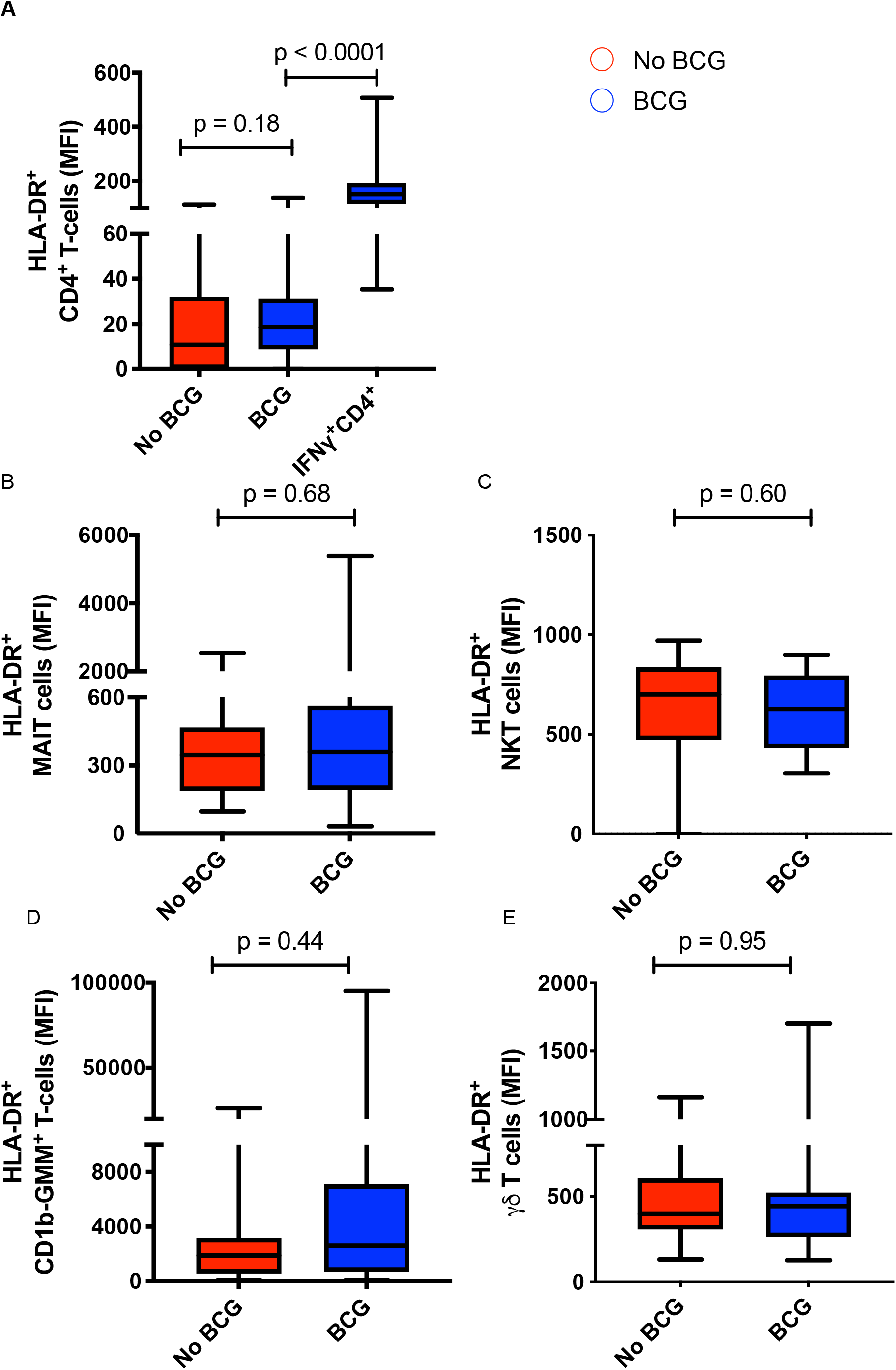
Assessment of BCG-associated T cell activation of the various cell subsets in BCG-vaccinated or unvaccinated infants. (A) T cell activation, measured by expression levels of HLA-DR, by total CD4^+^ T cells in BCG-vaccinated or unvaccinated infants or by IFN-γ-expressing, BCG-reactive CD4^+^ T cells in BCG-vaccinated infants (IFN-γ-expressing CD4^+^ T cells in unvaccinated infants were too infrequent to quantify HLA-DR expression). T cell activation of (B) MR1-5-OP-RU tetramer^+^ MAIT cells, (C) CD1d-PBS57 tetramer^+^ NKT cells, CD1b-GMM tetramer^+^ T cells or (E) γδ T cells. CD1b-GMM tetramer^+^ CD4^+^TRAV1-2^+^ GEM cells were too infrequent to reliably quantify HLA-DR expression. MFI, median fluorescence intensity. *P* values were computed by Mann-Whitney U test and p < 0.05 was considered statistically significant.

**Figure 4.**
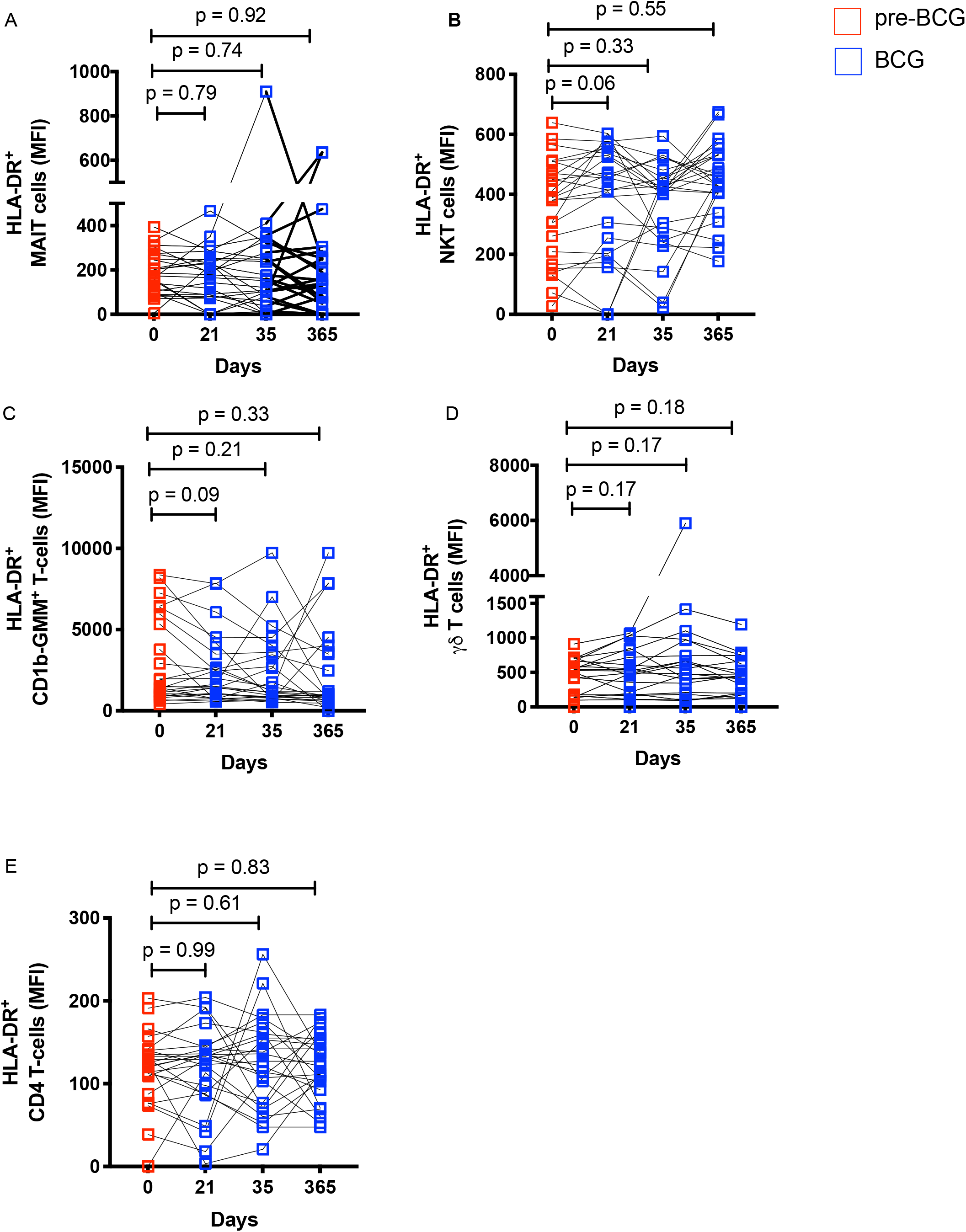
Longitudinal analysis of T cell activation profile of the various cell subsets in adults before and after BCG revaccination. T cell activation, measured by expression levels of HLA-DR, on MR1-5-OP-RU tetramer^+^ MAIT cells, CD1d-PBS57 tetramer^+^ NKT cells, CD1b-GMM tetramer^+^ T cells, γδ T cells or total CD4^+^ T cells. CD1b-GMM tetramer^+^ CD4^+^TRAV1-2^+^ GEM cells were too infrequent to reliably quantify HLA-DR expression. MFI, median fluorescence intensity. *P* values were computed by Wilcoxon signed-rank test and p < 0.05 was considered statistically significant.

### BCG vaccination modulates a novel CD26^+^CD161^+^ CD4 T cell subset in infants

We also sought to evaluate functional characteristics of DURT subsets. In a subset of the infant cohort, we quantified IFN-γ-expressing BCG-reactive γδ and phenotypically defined MAIT cells using a 12-hour whole blood intracellular cytokine-staining (WB-ICS) assay. Consistent with the higher overall frequencies of γδ T cells in BCG-vaccinated infants (Fig 2E), frequencies of BCG-reactive, IFN-γ-expressing γδ T cells tended to be somewhat higher in BCG vaccinated than unvaccinated infants (Fig.5A-B), although at p = 0.068 this did not reach the traditional level of significance of p < 0.05. In addition, frequencies of BCG-reactive IFN-γ-expressing CD26^+^CD161^+^ CD3^+^ T cells, which phenotypically meet the definition for MAIT cells (30), were significantly elevated (p<0.0045) in BCG vaccinated compared to BCG naïve infants (Fig. 5C-D). Further phenotypic characterization revealed that this result was driven by an IFN-γ-expressing, BCG-reactive CD4-positive CD26^+^CD161^+^ T cell subset, and not the typical CD8^+^ subset that characterizes MAIT cells (Fig. 5E-F). These BCG-reactive CD4^+^CD26^+^CD161^+^ T cells also did not express TRAV1-2, the canonical TCRα variable gene associated with MAIT cells (Fig. 5G). Unfortunately, the MR1 tetramer was not included in the flow cytometry panel used to analyze stimulated whole blood, and therefore we could not determine whether these cells were MR1-restricted. In summary, these data indicate that neonatal BCG vaccination modulates a functional subset of CD4^+^ T cells with a CD26^+^CD161^+^TRAV1-2^−^ phenotype.

**Figure 5.**
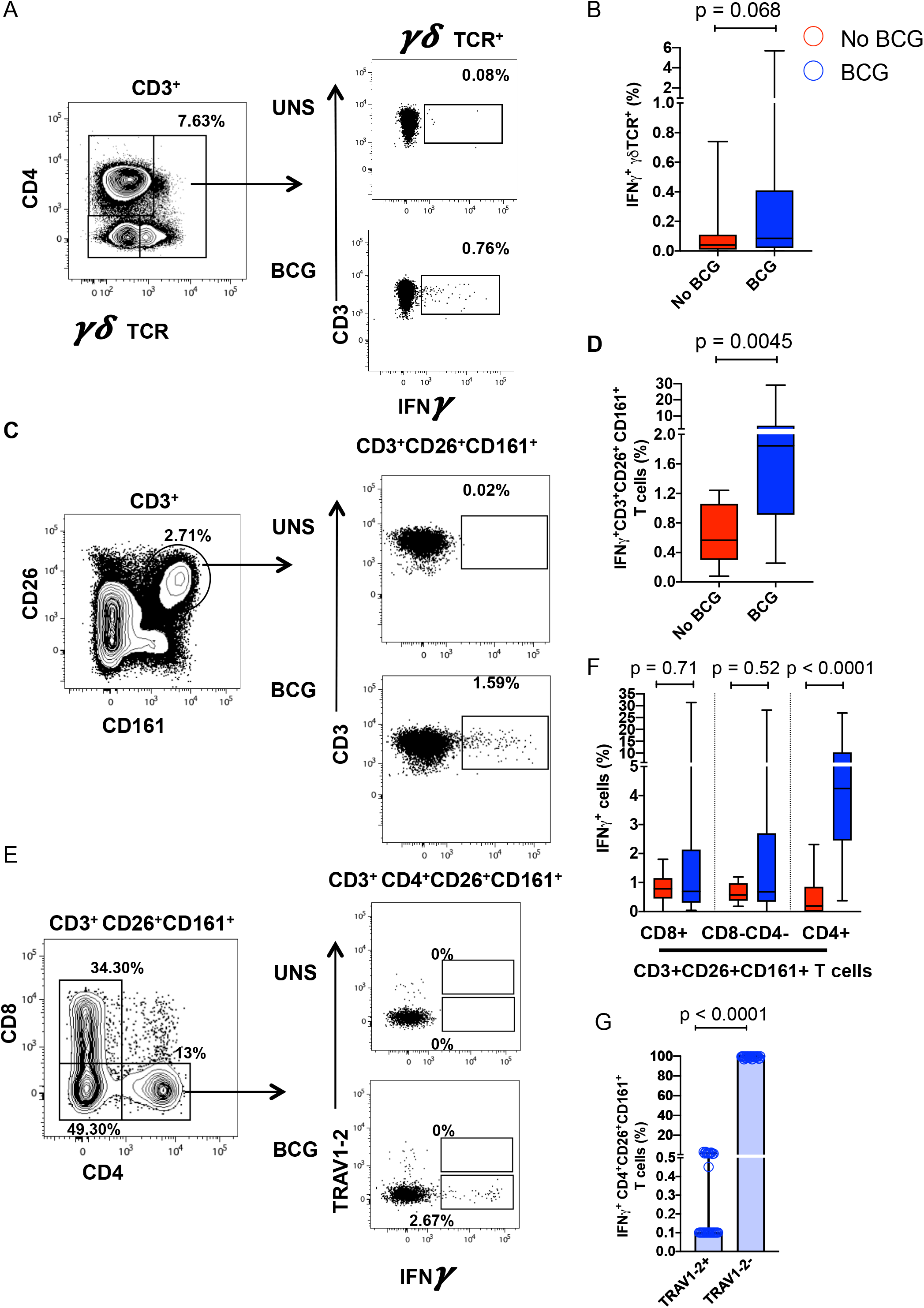
Unconventional IFN-y-expressing T cells elicited by infant BCG vaccination. (A) Representative flow cytometry plots depicting γδ TCR expressing CD3^+^ lymphocytes (left) and IFN-γ expression in unstimulated and BCG-stimulated γδ T cells (right). (B) Frequencies of BCG-reactive γδ T cells expressing IFN-γ in BCG-vaccinated (blue) and unvaccinated (red) infants. (C) Representative flow cytometry plots of (left) CD26 and CD161 expression by CD3^+^ lymphocytes to identify CD26^+^CD161^+^ T cells producing IFN-γ in unstimulated and BCG-stimulated infant blood samples (right). (D) Frequencies of BCG-reactive IFN-y-expressing CD3^+^CD26^+^CD161^+^ T cells in BCG-vaccinated (blue) and unvaccinated (red) infants. (E) Representative flow cytometry plot depicting CD8 and CD4 staining and gating in CD3^+^CD26^+^CD161^+^ T cells (left). Plot depicting TRAV1-2 and IFN-γ staining among CD4+CD26+CD161+ T cells (right). (F) Proportions of BCG-reactive IFN-γ+CD3^+^CD26^+^CD161^+^ T cells that are CD8+, double negative for CD8 and CD4 or that are CD4+ in BCG-vaccinated (blue) and unvaccinated (red) infants. (G) Proportions of BCG-reactive IFN-γ+CD4^+^CD26^+^CD161^+^ T cells that express TRAV1-2 in in BCG-vaccinated infants. Horizontal lines show medians, boxes the interquartile range and whiskers the range. *P* values were calculated by Mann-Whitney U-test and p < 0.05 was considered statistically significant.

### T cell memory profiles of DURT subsets differ from conventional CD4^+^ T cells

DURT cells are known to display immediate effector functions, such as secretion of inflammatory cytokines and cytotoxic molecules (31). By comparison, conventional MHC-restricted T cells fully develop effector functions only after antigen-induced priming and differentiation into memory and effector cells. To investigate possible effects of BCG on differentiation and memory marker expression, we assessed proportions of DURT cells expressing CCR7 and/or CD45RA.

In both infants and adults, MAIT and NKT cells predominantly expressed a CCR7^−^CD45RA^−^ phenotype at all time points, consistent with effector memory T cells (Fig. 6A-B). Moreover, this phenotype was not modulated following BCG vaccination in either study group. This “effector-like” phenotype was as prominent in infants as in the adult population, suggesting early development of this phenotype of MAIT and NKT cells in infants and little or no changes beyond early infancy. Longitudinal analysis of memory profiles in the adult cohort before and after BCG-revaccination suggested that BCG does not modulate CCR7 or CD45RA expression up to one year post-vaccination (Suppl. Figure 1). γδ T cells, on the other hand, exhibited a more mixed phenotype comprising CCR7^−^CD45RA^−^ (consistent with effector memory) and CCR7^−^ CD45RA^+^ (terminally differentiated effector) T cells (Fig. 6C). As expected from previous work, bulk CD4^+^ T cells in infants were comprised predominantly of naïve (CCR7^+^CD45RA^+^) and some central memory (CCR7^+^CD45RA^−^) T cells, whereas adults had comparatively lower proportions of naïve and higher proportions of central memory and effector memory CD4^+^ T cells (Fig. 6D).

**Figure 6.**
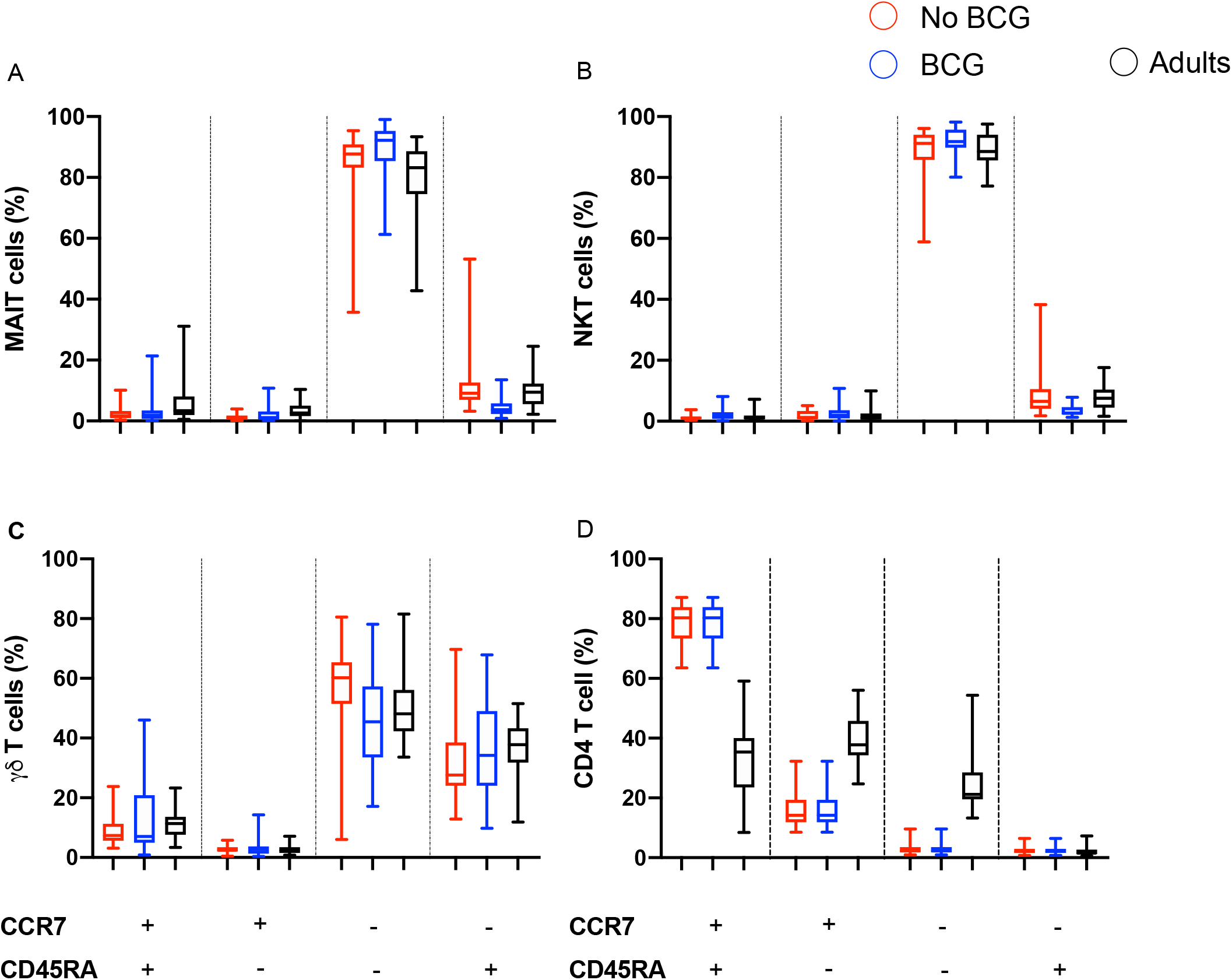
T cell memory phenotypes of DURT cell subsets or CD4 T cells in infants or adults. T cell memory profiles, measured by CCR7 and CD45RA co-expression patterns, on (A) MR1-5-OP-RU tetramer^+^ MAIT cells, (B) CD1d-PBS57 tetramer^+^ NKT cells, (C) γδ T cells or (D) total CD4 T cells in BCG-vaccinated (blue) and unvaccinated (red) infants or in adults (black) 5 weeks after BCG re-vaccination. Relative proportions of cells that fall into each of the possible combinations of CCR7 and CD45RA are represented as percentages. Boxes represent IQR, horizontal lines represent medians and whiskers represent the range. Longitudinal T cell memory profiles in adults are shown in Supplementary Figure 1.

## Discussion

DURT cells and their roles in protective and pathogenic immune responses have received significant attention in recent years (32). Because these cell populations are not donor restricted, are relatively abundant and are intrinsically poised for rapid effector function, there is growing interest in whether DURT cells can be harnessed to improve efficacy of vaccination against organisms that express ligands for DURT TCRs (1). We characterized frequencies, phenotypic and functional characteristics of DURT cell populations in the peripheral blood following BCG vaccination to explore their potential as possible immunological targets for TB vaccination. Evidence that γδ T cells, MAIT, NKT and GEM T cells can respond to mycobacteria (11, 15, 33–36) provides a strong rationale to investigate if BCG vaccination can modulate these T cell populations.

A key finding was that frequencies of γδ T cells, as well as frequencies of IFN-γ-expressing γδ T cells detected by intracellular cytokine staining after *in vitro* BCG stimulation, were increased in BCG-vaccinated compared with unvaccinated infants. However, BCG revaccination of adults did not result in significant changes in γδ T cell frequencies. Our results are consistent with data from non-human primates, in which a rapid and robust expansion of γδ T cells has been observed after BCG vaccination or M.tb-infection (18). In cattle, γδ T cells have been shown to accumulate in the early phases of *M. bovis* infection, but quickly dissipate in peripheral blood upon the arrival of other cells. Moreover, circulating γδ T cells from these animals produced increased amounts of IFN-γ and CCL2, and expressed high amounts of cytolytic molecules and lysed BCG-infected target cells with greater efficiency compared to cells from uninfected animals (37). Our findings are thus consistent with the previously reported role of γδ T cells as early responders to BCG vaccination, and may contribute to or enhance the ensuing immune response by recruiting or activating other key immune and effector response players. However, these results are at odds with findings of another recent study of γδ T cell development in early life. Frequencies and differentiation profiles of γδ T cells, assessed in cord blood and at 10 weeks of age, revealed a rapid expansion of Vγ9Vδ2 T cells with cytotoxic phenotypes after birth (38). Notably, this early and robust change in T cell responses was not attributable to newborn BCG vaccination, since Vγ9Vδ2 T cell responses between BCG-vaccinated and unvaccinated infants were not different. The data suggest a possible role of environmental phosphoantigen exposure in the priming of these cells in early life (38). The discrepancy between the BCG-attributable effects on γδ T cells between this study and our present one most likely relate to the different subsets of γδ T cells detected in each study. Although Vγ9Vδ2 are the main γδ T cell subset, in this study we measured the total γδ T cell population, and we cannot exclude that other minor subsets may have contributed to the differences observed here.

We found that infants generally had lower peripheral blood frequencies of MAIT, NKT, GMM-specific and GEM T cells compared to adults, a finding consistent with our previous study (39). By contrast, frequencies of γδ T cells were not different between infants and adults. As expected, BCG vaccination markedly induced conventional, IFN-γ-expressing CD4 T cell responses. However, we show that frequencies of MAIT, NKT, GMM-specific and GEM T cell populations were not modulated by intradermal BCG administration in either infants or adults. BCG vaccination also did not affect DURT cell activation or expression of the T cell memory markers CCR7 and CD45RA on DURT populations. The latter result is not surprising given that the DURT subsets exhibited very dominant CCR7^−^ CD45RA^−^ effector-like phenotypes in infants and adults, consistent with the well-described intrinsic effector function of these innate-like T cell populations (1).

The finding that BCG vaccination did not modulate DURT subsets can be explained by the well-described intrinsic innate-like and poor proliferative capacity generally attributed to DURT cells, which prescribe that these cells do not adapt to antigenic stimulation in the same manner as conventional, adaptive CD4 and CD8 T cells do. A recent study illustrated this elegantly by employing transcriptomic analyses of conventional T cells, MAIT cells, iNKT cells, γδ T cells and NK cells to investigate the apparent trade-off between potential for cellular proliferation and rapid effector function (40). The analyses revealed that these cell subsets can be ranked according to their “innateness” with conventional, adaptive CD4 T cells the least innate and NK cells the most, and that this innateness was characterized by pre-formed mRNA encoding effector molecules, while impaired proliferation was marked by decreased baseline expression of ribosomal genes. We acknowledge the possibility that the timing of our analyses, performed at 9 weeks after neonatal BCG vaccination and 3, 5 weeks and 1 year after adult BCG revaccination, may have missed transient DURT cell responses to BCG vaccination. In a controlled human Salmonella challenge model, MAIT cells were shown to be activated at the peak of infection (day 10) and this activation state was maintained even after antibiotic treatment (41). By contrast, there was an early decrease in MAIT frequencies after bacterial challenge (day 8), which recovered after antibiotic treatment (day 28) (41). Greene *et al*. (42) also reported significant, but transient activation of MAIT cells in peripheral blood of rhesus macaques in response to BCG, which peaked around 21 days after vaccination. Notably however, no significant modulation of MAIT cell frequencies in the peripheral blood was observed, highlighting that clonal expansion may be subtle and easy to miss. We also note that the adult BCG revaccinated cohort enrolled individuals with tuberculin skin test (TST) indurations exceeding 15mm, consistent with prior M.tb infection. The high levels of baseline immune sensitization in these individuals may have led to a masking effect such that subtle changes in BCG-induced immune responses were not detected. Future studies should also include TST or IGRA-negative individuals which are difficult to enrol in a high TB incidence setting like South Africa, to allow investigation of the effects of prior sensitization on vaccine-induced DURT responses. Our findings do not preclude that BCG vaccination may shape other qualitative aspects of DURT cells, such as their homing capacity, redistribution of the clonal TCR repertoire and/or proliferative potential.

Neonatal BCG vaccination was associated with a marked increase of an interesting and novel subset of BCG-reactive, IFN-γ-expressing CD26^+^CD161^+^ CD4 T cells. Further phenotypic characterization of this subset revealed that these cells did not express the canonical MAIT TCR alpha variable gene segment, TRAV1-2. MAIT cells have traditionally been phenotypically identified as TRAV1-2^+^CD161^+^ or CD26^+^CD161^+^ CD8^+^ cells, and most studies have focused exclusively on adults. We recently showed that MR1-5-OP-RU tetramer-positive MAIT cells in neonates and infants do also include CD4^+^ and TRAV1-2-negative T cells (39). In addition, in a TCR transgenic MAIT cell murine model, CD4^+^ MAIT cells were associated with pulmonary protective immunity, recruited into the lung in response to M.tb aerosol challenge (43). However, these BCG-reactive, IFN-γ-expressing CD26^+^CD161^+^ CD4 T cells could also be a subset of conventional, activated CD4 T cells or a novel DURT subset that are induced by BCG vaccination. Interestingly, these cells share characteristics with a recently described CD4^+^CD26^+^CD161^+^CCR6^+^ cell subset that expresses IL-17 and IL-22 and was preferentially enriched in a Peruvian cohort of TB non-progressors (44). These findings warrant further clinical and mechanistic investigation of these cells in clinical studies and animal models.

In conclusion, our study suggests that intradermal BCG vaccination does not modulate MAIT, NKT, GMM-specific and GEM T cell frequencies. However, newborn BCG vaccination was associated with increased frequencies of γδ T cells and a trend towards their enhanced production of IFNγ in response to BCG. More studies are required to understand the full potential of DURT cells for TB vaccination and whether the modulation of γδ T cells is durable. Given their immediate effector properties, the monoclonal or oligoclonal use of a TCR, and their role in regulating other key immune cell subsets and functions, future research should also explore their potential adjuvant effects.

## Methods

### Study participants

All participants were enrolled at the South African Tuberculosis Vaccine Initiative Field site in Worcester near Cape Town, South Africa. We collected blood from two cohorts of participants who were vaccinated with BCG.

The first cohort comprised two groups of healthy, 9-week-old infants. In one group (BCG vaccinated), infants received BCG at birth as is routine according to the Expanded Program on Immunization in South Africa. In the second group (BCG naïve), BCG vaccination was delayed until after blood collection at 9 weeks, for whole blood stimulation and PBMC cryopreservation. Infants who were born preterm (<37 weeks of gestation), with a low birth weight (<2.5 kg), had congenital malformations or perinatal complications, were in close contact with someone with TB disease or had suspected TB, received isoniazid preventive therapy or immunosuppressant therapy, or had any acute or chronic disease were excluded.

The second cohort, the TBRU adult BCG revaccination trial (28, 29), comprised healthy tuberculin skin test (TST)-positive, HIV-negative adults who received at least 6 months of isoniazid preventive therapy before or 7 months after BCG revaccination (NCT01119521). Peripheral blood was collected before revaccination and on days 21, 35, and 365 post-vaccination and peripheral blood mononuclear cells (PBMCs) cryopreserved.

### Ethics approvals

The studies were conducted in accordance with Good Clinical Practice (GCP) and guidelines set out by the World Medical Association’s Declaration of Helsinki. The studies and all procedures were reviewed and approved by the Human Research Ethics Committee (HREC) of the University of Cape Town as follows: BCG revaccination trial (Ref. 387/2008), infants vaccinated at birth (Ref. 126/2006) and infants with delayed BCG vaccination (Ref. 177/2011). The BCG revaccination study was also reviewed and approved by the University Hospital Cleveland Medical Center Institutional Review Board. Written informed consent was obtained from all adult participants and from a parent or legal guardian of infants before enrollment.

### Flow cytometry assays

#### PBMC staining

Cryopreserved PBMCs were thawed in a 37°C water bath, washed in PBS, and then stained with LIVE/DEAD dye according to the manufacturer’s instructions. Thereafter, PBMCs were stained with the fluorochrome-conjugated tetramers (CD1b-GMM (45), CD1d-PBS57, MR1-5-OP-RU, and their respective controls) for 45-60 minutes at room temperature, followed by CCR7 staining at 37°C for 30 minutes, and thereafter stained for other phenotypic markers (antibody panels described in Supplemental Table 1A) at 4°C for another 30 minutes.

#### Whole Blood intracellular cytokine staining (WB-ICS) assay and staining

Heparinized blood was processed, within 75 minutes of blood collection, using a standardized 12h WB-ICS assay protocol (46). Briefly, blood was stimulated with BCG Vaccine SSI (Biovac, Cape Town, South Africa) reconstituted with RPMI (final concentration 1.2 × 10^6^ CFU/ml), PHA (Sigma-Aldrich; positive control at 5 *μ*g/ml) or RPMI (negative control). For all stimulation conditions, co-stimulants anti-CD28 and anti-CD49d (BD Biosciences; San Diego, USA) were added at 0.25 *μ*g/ml. Blood was stimulated for 7 hours at 37°C, after which Brefeldin-A (Sigma-Aldrich) was added at a concentration of 10 *μ*g/ml for the remaining 5 hours of stimulation. At the end of the stimulation, 2mM of EDTA (Sigma-Aldrich) was added, red blood cells were lysed using 1:10 FACS Lysing solution (BD Biosciences) and fixed white blood cells were cryopreserved in liquid nitrogen. Cryopreserved fixed white blood cells from infant participants were thawed, washed in PBS, permeabilized in Perm/Wash buffer (BD Biosciences) and stained with antibody panels (Supplemental Table 2, 3) for 30 minutes at 4°C.

#### Flow cytometry

Stained samples were acquired on a BD-LSR-II flow cytometer configured with 4 lasers: Solid state Blue (488nm; 100mW; 3 detectors), Solid state Violet (405nm; 25nW; 8 detectors), HeNe gas Red (635nm; 70mW; 3 detectors), and Diode-pumped Coherent Compass (532nm; 150mW; 8 detectors). We used mouse κ chain BD CompBeads stained with each individual antibody conjugate (for tetramer reagents, fluorochrome-matched antibody conjugates were used) to compensate all parameters. Samples were acquired with optimal photomultiplier tube voltages calibrated daily using targets for SPHERO Rainbow Fluorescent Particles (Spherotech, Inc.).

## Data analysis

Flow cytometric data were analyzed using FlowJo (version 10.5.3). Statistical analyses were performed in GraphPad Prism (version 7). Mann-Whitney U test was used to compare responses between vaccinated and non-vaccinated infant groups, whereas a Wilcoxon signed-rank test was used to compare longitudinal responses within the adult population. p-values < 0.05 were considered statistically significant.

## Acknowledgments

We would like to thank the participants who gave their time and dedication to this study. This study was initiated and designed by members of the Collaboration for TB Vaccine Discovery (CVTD), DURT Research Community, with support from the Bill and Melinda Gates Foundation. The study was funded by a research grant from Aeras. The delayed BCG infant study was funded by NIH R01 grant AI087915 and the adult BCG revaccination study was funded by NIH grant NO1-AI70022. CD1b tetramers, ligands and their validation was supported by R01 AI049313. Anele Gela was supported by postdoctoral fellowships from the Claude Leon Foundation and the Harry Crossley Foundation. Melissa Murphy was supported by a Masters and Doctoral Innovation Scholarship from the National Research Foundation. The funders had no role in study design, data collection and analysis, decision to publish, or preparation of the manuscript. The MR1 tetramer technology was developed jointly by Dr. James McCluskey, Dr. Jamie Rossjohn, and Dr. David Fairlie, and the material was produced by the NIH Tetramer Core Facility as permitted to be distributed by the University of Melbourne.

## Contributions

AG, DFH, WAH, WHB, JLJ, DFH, SAJ, THMO, SS, DBM, DL, MH, CS, EN and TJS designed the experiments or clinical studies. WAH, HB, and MH performed clinical investigations. AG, MM, KH and SS processed samples, performed assays and/or analyzed data. AG, MM, KH, SS, EN and TJS interpreted the data. AG, MM, CS, EN and TJS wrote the manuscript. All authors reviewed and approved the manuscript.

## Supplementary Information

Supplementary information is available in the online version of this publication.

---

**Suppl. Figure 1.**
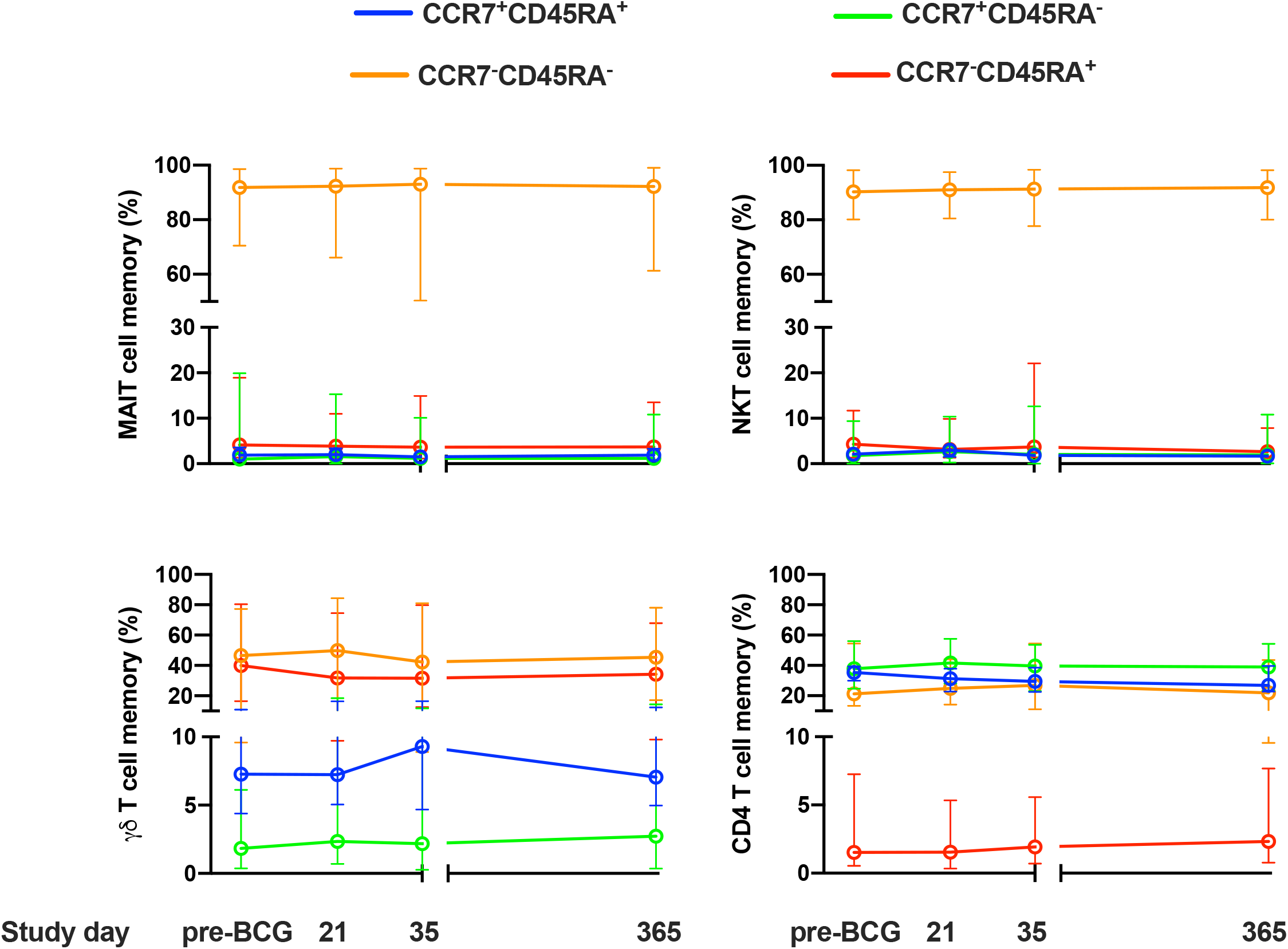

